# CRISPR interference at the *FAAH-OUT* genomic region reduces *FAAH* expression

**DOI:** 10.1101/633396

**Authors:** Hajar Mikaeili, Charlix Yeung, Abdella M Habib, John N Wood, Andrei L Okorokov, James J Cox

## Abstract

*FAAH-OUT* encodes a putative long non-coding RNA which is located next to the *FAAH* gene on human chromosome 1. Recently an ~8 kb microdeletion, that removes upstream regulatory elements and the first 2 exons of *FAAH-OUT*, was reported in a pain insensitive patient (PFS) with additional clinical symptoms including a happy, non-anxious disposition, fast wound healing, fear and memory deficits, and significant post-operative nausea and vomiting induced by morphine. PFS also carries a hypomorphic SNP in *FAAH* that significantly reduces the activity of the encoded fatty-acid amide hydrolase enzyme. The *FAAH* and *FAAH-OUT* mutations identified in PFS result in elevated levels of anandamide (AEA), palmitoylethanolamide (PEA) and oleoylethanolamine (OEA) measured in peripheral blood. These bioactive lipids, which are normally degraded by FAAH, have diverse biological functions including known roles in pain pathways.

Here we report the first mechanistic insights into how the *FAAH-OUT* microdeletion affects *FAAH* function. Gene editing in a human cell line to mimic the *FAAH-OUT* microdeletion observed in PFS results in reduced expression of *FAAH*. Furthermore, CRISPRi experiments targeting the promoter region of *FAAH-OUT* or a short highly evolutionarily conserved element in the first intron of *FAAH-OUT* also result in reduced expression of *FAAH*. These experiments confirm the importance of *FAAH-OUT* and specific genomic elements within the ~8 kb microdeleted sequence to normal *FAAH* expression. Our results also highlight the potential of CRISPRi and gene editing strategies that target the *FAAH-OUT* region for the development of novel *FAAH*-based analgesic and anxiolytic therapies.

---

Millions of people worldwide are living in chronic pain^1^. This pain is often poorly treated and the over-prescription of opioid-based drugs has contributed to an opioid epidemic that is causing significant morbidity and mortality^2^. New pain-killing and non-opioid based medications are hence urgently needed. A powerful way to identify novel human-validated analgesic drug targets is to study rare individuals with intact damage-sensing neurons that present with a congenital pain insensitive phenotype^3,4^. For example, the study of pain-free individuals from three consanguineous families from northern Pakistan led to the identification of recessive mutations in *SCN9A* as a cause of painlessness^5^. *SCN9A*, which encodes the voltage-gated sodium channel, Na_v_1.7, is highly expressed in damage-sensing neurons and is now a major drug target in the pharmaceutical industry with several pain clinical trials in progress^6^. Likewise, the study of an Italian family in which six affected individuals displayed a pain insensitive phenotype led to the identification of a point mutation in the *ZFHX2* transcriptional regulator, a gene also with enriched expression in peripheral sensory neurons^7^.

Recently a new pain insensitivity disorder was reported from studying a female patient (PFS) who carried both a hypomorphic SNP in the fatty-acid amide hydrolase (*FAAH*) gene and a microdeletion downstream of *FAAH* in a novel gene called *FAAH-OUT*^8^. This patient, in addition to being pain insensitive, also presented with additional clinical symptoms including a happy, non-anxious disposition, enhanced wound healing, reduced stress and fear symptoms and mild memory deficits. FAAH is a major catabolic enzyme for a range of bioactive lipids called fatty-acid amides (FAAs) with substrates including anandamide (AEA), palmitoylethanolamide (PEA), oleoylethanolamine (OEA) and N-acyltaurines^9–11^. The phenotype observed in PFS likely results from elevated levels of these bioactive lipids due to reduced activity of FAAH. Indeed, mass spectrometry analysis of peripheral blood derived from PFS showed significantly raised levels of AEA, OEA and PEA^8^.

How does the ~8 kb microdeletion that is distinct from and begins ~5 kb downstream of the 3’ end of the currently annotated footprint of the *FAAH* gene disrupt its function? Potential mechanisms include (1) the microdeleted genomic sequence contains important regulatory elements needed for normal *FAAH* expression (e.g. an enhancer); (2) the *FAAH-OUT* transcript has an epigenetic/transcriptional role in regulating *FAAH* expression; (3) the *FAAH-OUT* transcript normally acts as a decoy for microRNAs that also target *FAAH* mRNA; (4) the *FAAH-OUT* gene encodes a short functional protein; (5) *FAAH* mis-splices into *FAAH-OUT* resulting in an aberrant, non-functional transcript. Here we show by gene editing in human cells that removing the ~8 kb region that is deleted in PFS results in reduced expression of *FAAH*. Furthermore, targeting dSaCas9-KRAB to the *FAAH-OUT* promoter region or to a short evolutionarily conserved ‘FAAH-AMP’ element within the first intron of *FAAH-OUT*, also leads to reduced *FAAH* mRNA levels. These CRISPRi experiments highlight specific regions within the *FAAH-OUT* genomic region that may be targeted as a gene therapy approach for treating pain.

## Methods

### CRISPR/Cas9 plasmids

Plasmids 61591^12^ and 106219^13^ (Addgene) were modified for the gene editing (SaCas9) and transcriptional repression (dSaCas9-KRAB) CRISPR experiments. For gene editing plasmid 61591, the CMV promoter was replaced with a shorter promoter sequence derived from the housekeeping *Eef1a1* gene and the bGH polyadenylation sequence was replaced with a shorter synthetic polyadenylation sequence. gBlocks gene fragments (IDT) were designed to contain a U6 promoter, guide sequence and modified guide scaffold^14^ with the design enabling two guide cassettes to be inserted into one plasmid by In-Fusion cloning (Takara). Guide sequences were designed using the CRISPOR tool^15^ and chosen to flank the microdeletion observed in patient PFS. Plasmid HMa included guides CCCAGTGAGTACGATGGCCAG and TTAGTGATATTGTTCCGTGGG. Plasmid HMb contained guides TCATGGCCTTTCCCCTTCTCA and GTCACTTGCAGTCTGATTAAG. The ‘empty vector’ control contained *Eef1a1*-promoter driven SaCas9 but no guide sequences.

For the transcriptional repression CRISPRi experiments the AgeI-EcoRI fragment from plasmid 106219 containing the dSaCas9-KRAB sequence was used to replace the SaCas9 sequence from plasmid 61591, to give a CMV driven dSaCas9-KRAB. Next, a gBlocks gene fragment (IDT) was designed to contain a synthetic poly(A) sequence, U6 promoter, guide sequence and modified guide scaffold^14^. This sequence was cloned into the EcoRI-NotI sites of the modified plasmid 61591 using In-Fusion cloning (Takara). The guide sequence ‘FOP1’ (AAAAGGTGAGGTCACGAGGCC) was located within a DNase hypersensitivity site approximately 323 bp upstream of the *FAAH-OUT* transcriptional start site. The guide sequence ‘FOC1’ (TGTTCCCATGCTTTGGAGTTC) was located at the highly evolutionarily conserved element located in the first intron of *FAAH-OUT*. The ‘empty vector’ control contained the CMV driven dSaCas9-KRAB but no guide sequence.

### Transfection of CRISPR/Cas9 plasmids into HEK293 cells

Lipofectamine 3000 (Invitrogen) was employed as a DNA carrier for transfection into human embryonic kidney 293 cells (ECACC) according to the manufacturer’s procedures. The HEK293 cells were cultured in Dulbecco’s modified Eagle’s medium (Life Technologies, Inc.) with 10% fetal bovine serum (Hyclone). Lipofectamine 3000 was diluted into Opti-MEM I Reduced Serum Medium (Life Technologies, Inc.). 5μg of plasmid DNA was first diluted into Opti-MEM and 10μl of P3000 reagent was added to the mixture. The DNA-liposome complex was prepared by adding diluted DNA into diluted Lipofectamine (ratio 1:1) and incubating the mixture at room temperature for 30 min. DNA-liposome mixture was added to 70% confluent HEK293 cells. After 24 hours of incubation at 37 °C, media was removed and the transfection steps were repeated. The cells were incubated at 37°C in a 5% CO_2_ incubator with 92-95% humidity for another 24 hours. To extract total RNA from cultured cells, medium was first aspirated off and cells were rinsed with ice-cold PBS. 1ml of Trizol^®^ was added directly to the cells and was incubated for 5 minutes at room temperature. Cell lysate was passed through a pipette up and down several times. RNA was extracted using PureLink™ RNA Micro Scale Kit (Invitrogen) according to the manufacturer’s procedures. Genomic DNA was isolated using the DNeasy Blood and Tissue kit (Qiagen) and used as template to confirm gene editing. Primers 5’TTAATGTCTGGAGTGATAACATGAC and 5’ACAACTTCTAATTAGTGTTAATGAC were used to amplify a ~463 bp band from HMa transfected cells and a ~598 bp band from HMb transfected cells. The size of the microdeletion induced by plasmids HMa and HMb was ~9017 bp and ~8882 bp respectively.

### TaqMan real-time PCR

HEK293 cell RNA (5 μg) (from the CRISPR/Cas9 experiments) was reverse transcribed using oligo d(T) and Superscript III first-strand synthesis system (Invitrogen) according to the manufacturer’s conditions. TaqMan real-time PCR was carried out using the following probes: *FAAH* (Hs01038660_m1), *FAAH-OUT* (Hs04275438_g1) and *Actin* (Hs01060665_g1). *FAAH* or *FAAH-OUT* expression was compared with that of *Actin* (*ACTB*) measured on the same sample in parallel on the same plate, giving a CT difference (ΔCT) for *ACTB* minus the test gene. Mean and standard errors were performed on the ΔCT data and converted to relative expression levels (2^ΔCT).

## Results

### Gene editing mimicking the *FAAH-OUT* microdeletion reduces *FAAH* expression

Patient PFS carries a 8,131bp microdeletion on chromosome 1 (hg19, chr1:46,884,415-46,892,545) that begins approximately 4.7kb downstream of the *FAAH* 3’ UTR (Figure 1)^8^. The microdeletion contains the first two exons and putative promoter region of *FAAH-OUT*, a novel long non-coding RNA with a similar tissue expression profile to *FAAH*. Gene editing experiments in human embryonic kidney cells transiently transfected for 48 hrs with an SaCas9 plasmid bearing 2 guide sequences that target sequences flanking the microdeletion showed the expected genomic deletion (Figure 2A). Next, total RNA was isolated from the HEK293 cells (a mixture of transfected and untransfected cells) and reverse transcribed into cDNA. Quantitative real-time PCR showed a significant reduction in both *FAAH-OUT* and *FAAH* mRNAs for cells transfected with each set of guide pairs (HMa and HMb) that flank the microdeletion, highlighting that *FAAH* expression is affected by the induced downstream deletion (Figure 2B).

**Figure 1:**
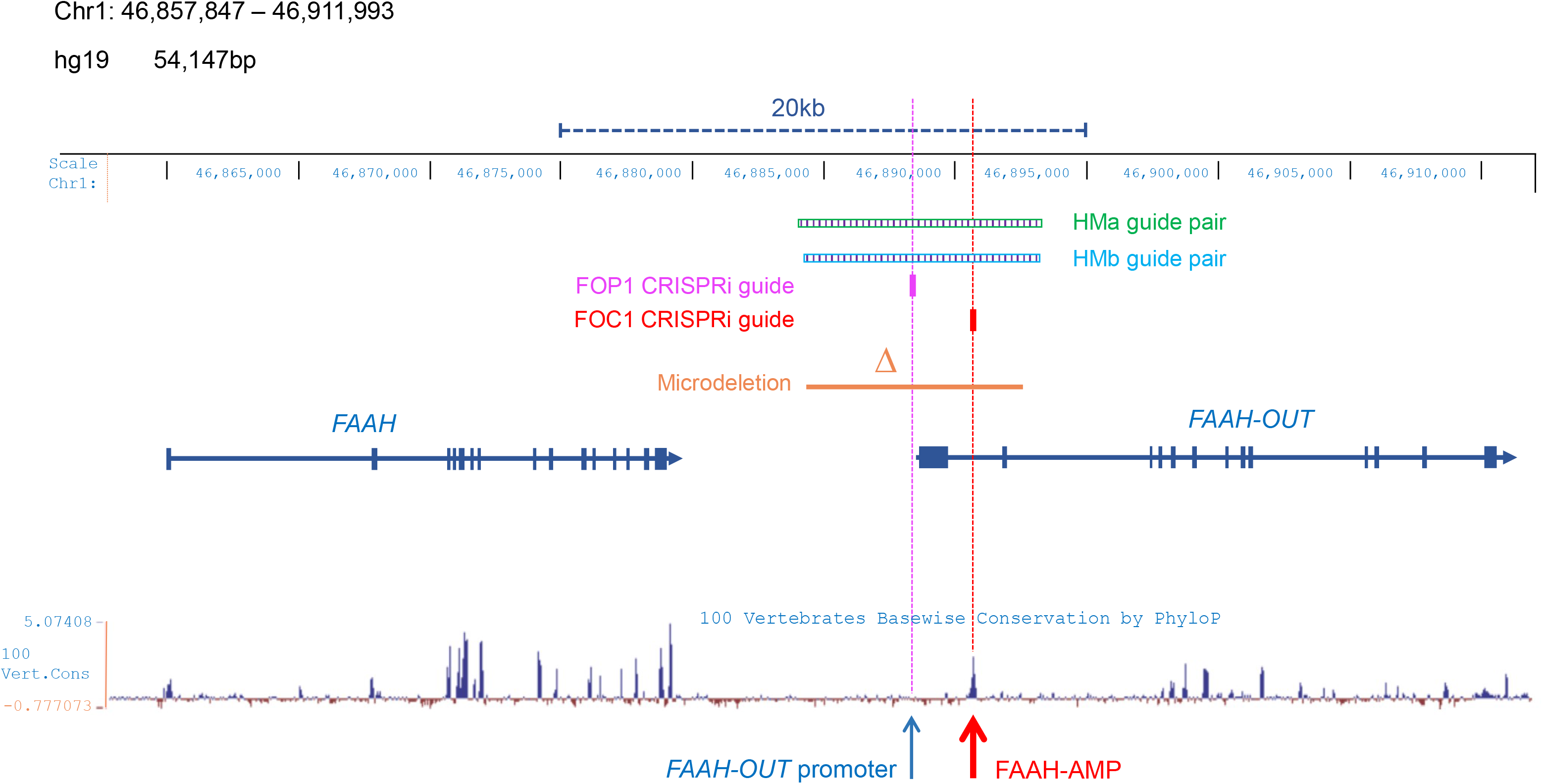
*FAAH* and *FAAH-OUT* genomic region. Map showing human chromosome 1 (46,857,847 – 46,911,993; build hg19). *FAAH* and *FAAH-OUT* genes are shown with exons denoted by blue boxes; the direction of transcription shown by arrows. The ~8 kb microdeletion identified in patient PFS is shown by the orange bar. Gene editing guide pairs HMa and HMb flank the microdeleted region. The CRISPRi guide FOP1 maps to the promoter region of *FAAH-OUT*; FOC1 maps to the first intron of *FAAH-OUT* and is located at the evolutionarily conserved ‘FAAH-AMP’ element (red arrow). The PhyloP basewise conservation track for 100 vertebrates from the UCSC genome browser shows regions of high conservation as peaks, with the majority of these mapping to gene exons in *FAAH* and *FAAH-OUT*.

**Figure 2:**
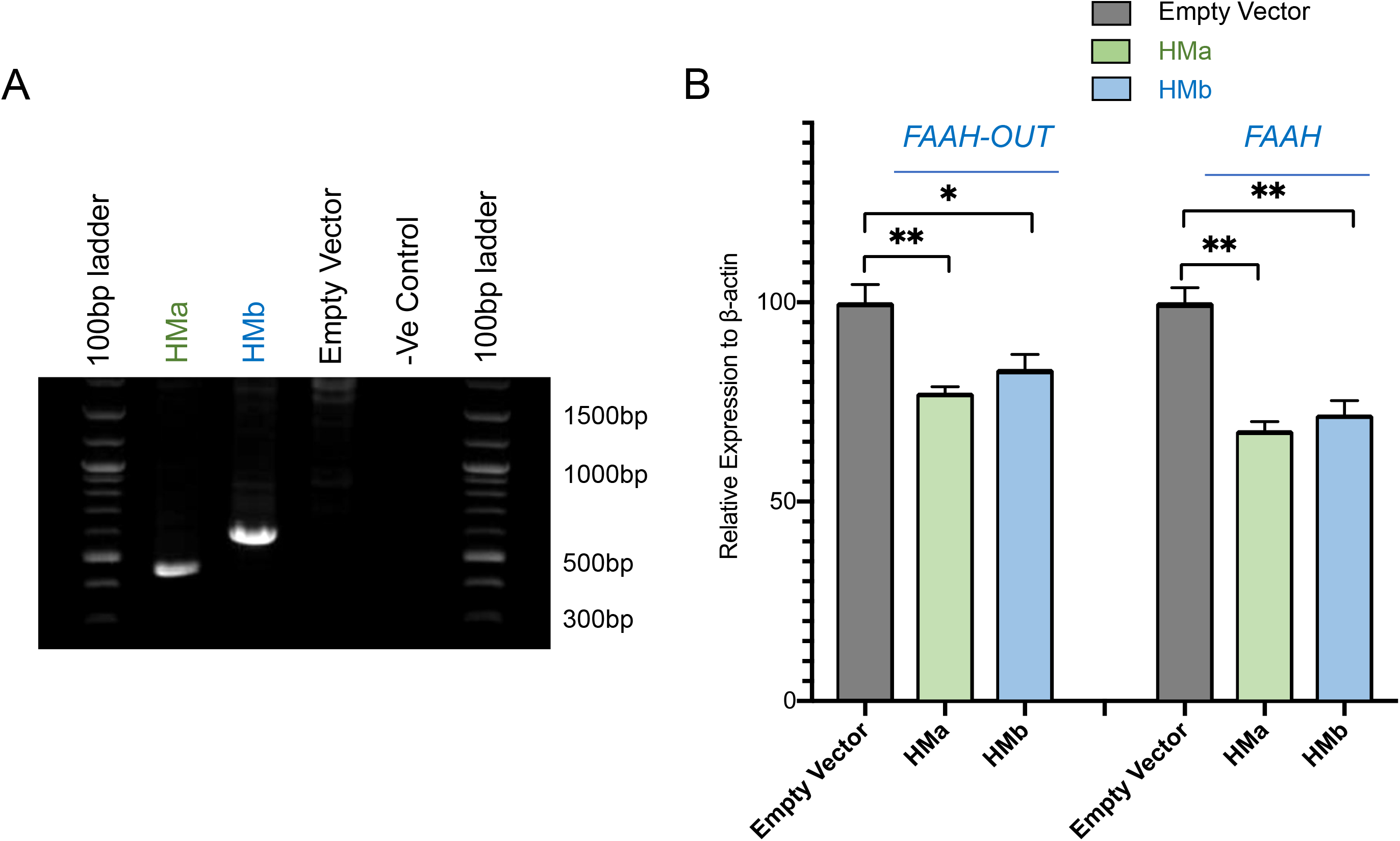
Gene editing mimicking the *FAAH-OUT* microdeletion reduces *FAAH* expression. **(A)** PCR reactions using primers that flank the gene editing HMa and HMb guide pairs; template genomic DNA isolated from dishes of HEK293 transiently transfected (48 hrs) with the SaCas9 plasmids. Gene editing is detected by a ~463 bp fragment amplified from HMa edited cells and a ~598 bp fragment from HMb edited cells. No band is observed from empty vector transfected cells indicating no editing at this locus (the large size of the unedited allele is beyond the capability of the DNA polymerase). **(B)** RT-qPCR analysis of both *FAAH-OUT* and *FAAH* mRNA levels following transient transfection with HMa or HMb SaCas9 plasmids. The microdeletion in *FAAH-OUT* results in a significant reduction in both *FAAH-OUT* and *FAAH* expression. *P* values were generated by t-test, *p<0.05; \*\**p*<0.01. Error bars = SEM

### CRISPRi at the *FAAH-OUT* promoter reduces both *FAAH-OUT* and *FAAH* expression

CRISPR interference (CRISPRi) enables gene repression through targeting of a nuclease-deficient form of Cas9 (dCas9) fused to a Krüppel-associated box (KRAB) repressor to specific genomic loci^16,17^. When localised to DNA, dCas9-KRAB recruits a heterochromatin-forming complex that causes histone methylation (H3K9 trimethylation) and deacetylation. This local modification of the epigenome can help to identify promoter and enhancer elements essential for gene expression^17,18^. A guide sequence (FOP1) was designed to recruit dSaCas9-KRAB to the putative promoter region of *FAAH-OUT* (Figure 1), approximately 300 bp upstream of the transcriptional start site previously identified by 5’ RACE^8^. Transient transfection of HEK293 cells with the FOP1-dSaCas9-KRAB plasmid followed by real-time qPCR showed a significant reduction in both *FAAH* and *FAAH-OUT* expression (Figure 3, bars in magenta) compared to empty vector transfected controls. This result indicates that transcription of the *FAAH-OUT* long non-coding RNA contributes to normal expression of *FAAH* and may be acting as an enhancer RNA, perhaps similar to how *lincRNA-Cox2* functions to regulate the upstream *Ptgs2* gene^19^.

**Figure 3:**
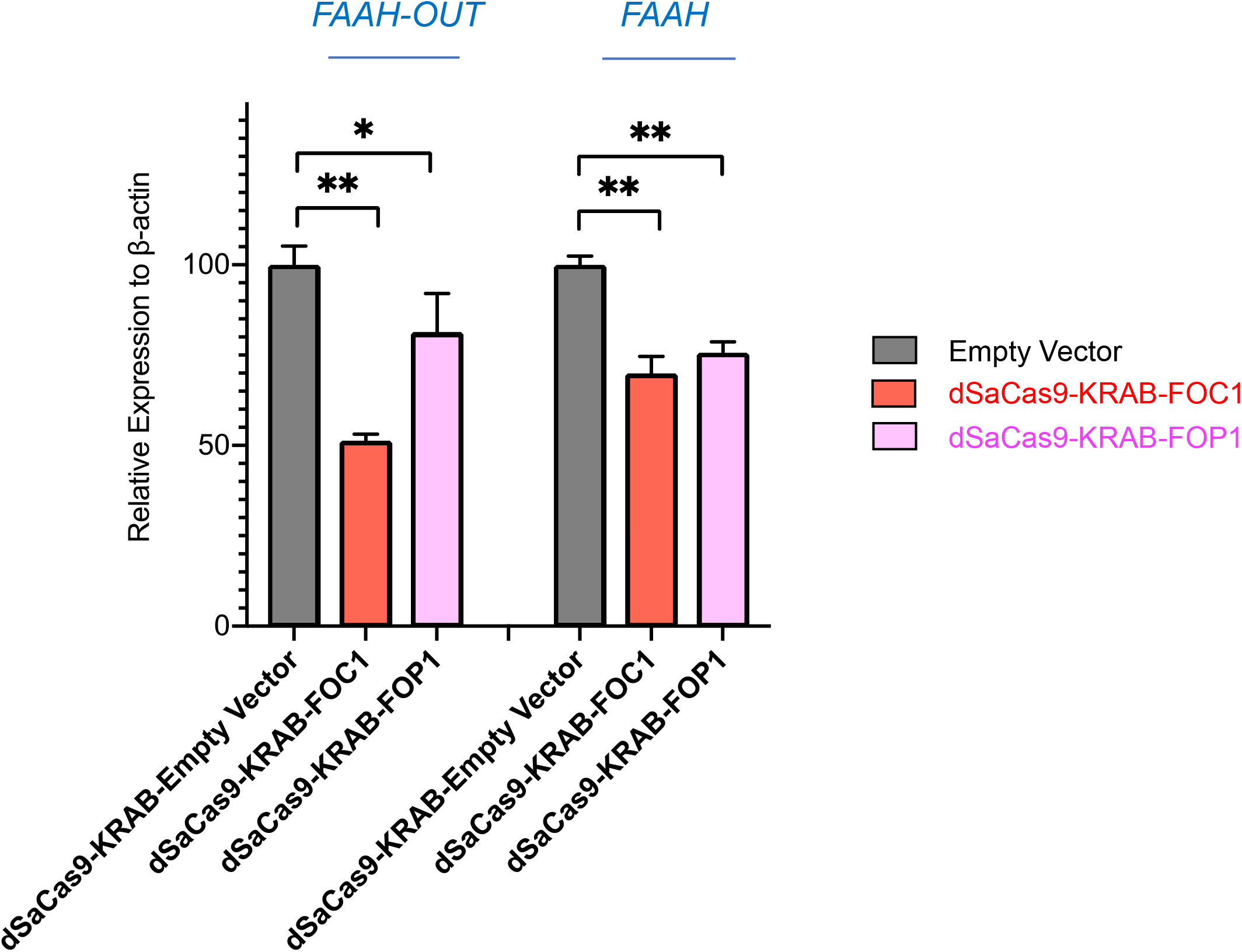
CRISPRi at the *FAAH-OUT* promoter and at a highly conserved ‘FAAH-AMP’ element reduces both *FAAH-OUT* and *FAAH* expression. dSaCas9-KRAB directed to the promoter of *FAAH-OUT* by guide ‘FOP1’ results in a reduction in both *FAAH-OUT* and *FAAH* expression (bars in magenta), whilst dSaCas9-KRAB directed to the FAAH-AMP evolutionarily conserved element in intron 1 of *FAAH-OUT* by guide ‘FOC1’ also results in a significant reduction in both *FAAH-OUT* and *FAAH* expression (bars in red) as measured by qPCR. *P* values were generated by t-test, \**p*<0.05; \*\**p*<0.01. Error bars = SEM

### CRISPRi at a highly conserved ‘FAAH-AMP’ element reduces *FAAH* and *FAAH-OUT* expression

Comparative genomic analyses across species can help to identify evolutionarily conserved sequences that may have important functions^20^. By analysing the PhyloP basewise conservation track for 100 vertebrates on the UCSC genome browser, a highly conserved element (denoted ‘FAAH-AMP’) was identified in the first intron of *FAAH-OUT* (Figure 1). We considered that this region may contain important regulatory sequences for *FAAH-OUT* and/or *FAAH*. To test whether this region may be acting as a critical enhancer element, a guide sequence (FOC1) was designed to recruit dSaCas9-KRAB to this highly conserved region (Figure 1). Transient transfection of HEK293 cells with the FOC1-dSaCas9-KRAB plasmid followed by real-time qPCR showed a significant reduction in both *FAAH* and *FAAH-OUT* expression (Figure 3, bars in red) compared to empty vector transfected controls. This result indicates that this putative FAAH-AMP enhancer element contributes to normal *FAAH* expression. Interestingly, the ‘HMR (human/mouse/rat) conserved transcription factor binding site’ track on the UCSC human genome browser shows several potential binding sites within the FAAH-AMP region, including those for S8, Chx10, Cart-1, LUN-1 and CUTL1 transcription factors.

## Conclusions

In summary, we provide the first mechanistic insights into how the microdeletion identified in patient PFS negatively affects *FAAH* expression. The ~8 kb microdeletion contains the upstream promoter region and first two exons of *FAAH-OUT* and also a highly evolutionarily conserved ‘FAAH-AMP’ element in the first intron that includes several potential transcription factor binding sites. Further work will help to understand precisely how *FAAH-OUT* may be functioning as an enhancer RNA and also clarify the importance of the FAAH-AMP element to normal *FAAH* expression.

## Acknowledgements

We gratefully acknowledge the support of the Medical Research Council (G1100340 and MR/R011737/1) and Wellcome (200183/Z/15/Z).

